# Inhibition of GFAT1 in lung cancer cells destabilizes PD-L1 protein and improves immune response

**DOI:** 10.1101/2020.11.17.385039

**Authors:** Wenshu Chen, Bryanna Saxton, Steven A. Belinsky

## Abstract

Immunotherapy using checkpoint blockers (antibodies) has been a major advance in recent years in the management of various types of solid cancers including lung cancer. One target of checkpoint blockers is programmed death ligand 1 (PD-L1) expressed by cancer cells, which engages programmed death 1 (PD-1) on T cells and Natural Killer (NK) cells resulting in suppression of their activation and cancer-killing function, respectively. Apart from antibodies, other clinically relevant agents that can inhibit PD-L1 are limited. PD-L1 protein stability depends on its glycosylation. Here we show that L-glutamine:D-fructose amidotransferase 1 (GFAT1) a rate-limiting enzyme of the hexosamine biosynthesis pathway (HBP) which produces uridine diphosphate-N-acetyl-β-glucosamine (UDP-GlcNAc), a precursor for glycosylation, is required for the stability of PD-L1 protein. Inhibition of GFAT1 activity markedly reduced interferon γ (IFNγ)-induced PD-L1 levels in various lung cancer cell lines. GFAT1 inhibition suppressed glycosylation of PD-L1 and accelerated its proteasomal degradation. Importantly, inhibition of GFAT1 in IFNγ-treated cancer cells enhanced the activation of T cells and the cancer-killing activity of NK cells. These findings support using GFAT1 inhibitors to manipulate PD-L1 protein level that could augment the efficacy of immunotherapy for lung cancer.

## Introduction

Cancer immunotherapy using immune checkpoint blockers reinvigorates dysfunctional cytotoxic T cells in cancer tissues to eliminate cancer cells. Current checkpoint blockers, i.e., monoclonal antibodies against cytotoxic T lymphocyte antigen 4 (CTLA-4), programmed death 1 (PD-1) and programmed death ligand 1 (PD-L1), disrupt the inhibitory signaling that limits T cell activity(1). Checkpoint blockers provide a vital option for treatment of many types of solid cancers including lung cancer(2,3). Despite the exciting clinical outcome, the therapeutic benefits have been limited to a minority of patients, and the response rate remains moderate and relapse is common. Understanding the mechanisms of resistance and the options to overcome them is critical to bringing immunotherapy with better efficacy to more patients.

Although a high level of PD-L1 in cancer cells is used as an indicator of a “hot” immune status that predicts likely response to PD-1/PD-L1 antibodies, the protein suppresses the activities of cytotoxic T cells and NK cells that express PD-1 to reduce their effectiveness against cancer, thus contributing to resistance to immune and conventional therapies(4–6). In general, a high level of PD-L1 in the cancer cell is prognostic for worse survival of patients with non-small cell lung cancer(7,8).

PD-L1 expression is controlled at transcriptional and posttranscriptional levels in cancer cells. The protein can be induced by various growth factors and cytokines(9). For example, IFNγ, released by T cells upon activation to attack cancer cells, strongly increases PD-L1 transcription (4,5). As a membrane protein, the level of PD-L1 protein in a cell is affected by posttranslational modification such as glycosylation, ubiquitination, trafficking, and recycling. Glycosylation of PD-L1 was recently shown to control its degradation via proteasomal degradation pathway(9).

Hexosamine biosynthesis pathway (HBP) uses glucose, glutamine, and nucleotide to produce uridine diphosphate-N-acetyl-glucosamine (UDP-GlcNAc), a building block for glycosylation of a variety of proteins(10). In this study, we show that GFAT1, the rate-limiting enzyme of HBP, regulates IFNγ-induced PD-L1 expression in lung cancer cells. Inhibition of GFAT1 reduced PD-L1 by accelerating proteasomal degradation of the protein. Importantly, inhibition of GFAT1 in cancer cells improves T cell response and killing of cancer cells by NK cells. Thus, our findings support GFAT1/HBP as a potential target for improving cancer immunotherapy.

## Materials and Methods

### Reagents and antibodies

Phorbol 12-myristate 13-acetate (PMA, P8139), ionomycin (I9657) and tunicamycin (T7765) were purchased from Sigma (St. Louis, MO, USA) and prepared in dimethyl sulfoxide (DMSO). Azaserine (A4142), 6-diazo-5-oxo-L-norleucine (DON, D2141), 3-methyladenine (M9281), and MTT [(3-(4,5-dimethylthiazol-2-yl)-2,5-diphenyltetrazolium bromide, M5655] were also Sigma products and prepared in distilled water or phosphate buffered saline. CB-839 (1439399-58-2) was obtained from Cayman Chemical (Ann Arbor, MI). VPS34 inhibitor (532628) was from Millipore (Billerica, MA). Recombinant human IFNγ (285-IF) was purchased from R & D Systems (Minneapolis, MN) and mouse IFNγ (BMS326) from eBioscience (Thermo Fisher Scientific, Waltham, MA). The following antibodies were used for Western blot: GFAT1 (ab125069), GFAT2 (ab190966), and RL2 (ab2739, recognizing O-GlcNAc) from Abcam (Cambridge, MA); human interferon gamma response factor 1 (IRF-1, sc-74530), O-GlcNAc transferase (OGT, sc-32921), and β-actin (sc-4778) from Santa Cruz Biotechnology (Santa Cruz, CA); human PD-L1 (AF156, goat) and mouse PD-L1 (AF1019, goat) from R & D systems; GFAT1 (5322), GFAT2 (6917) and PD-L1 (13684, rabbit) from Cell Signaling; β-actin (A2103) and β-tubulin (T6074) from Sigma.

### Cell culture

All cell lines were obtained from American Type Culture Collection (ATCC, Manassas, VA). Lung cancer cell lines (A549, H1975 and H2009, adenocarcinoma; SK-MES1 and SW900, squamous cell carcinoma) were authenticated by short tandem repeat (STR) DNA profiling using commercial service (Genetica Cell Line Testing, Burlington, NC) in 2017 (A549 and H1975) or 2018 (H2009, SK-MES-1 and SW900). Murine Lewis lung cancer cell line (LLC1) was purchased in 2016 and NK-92 Natural Killer cell line in 2020. Authenticated and recently purchased cell lines were expanded and frozen as stocks for subsequent recoveries and use. Jurkat T cell line was not authenticated. Cancer cell lines and Jurkat T cells were maintained in RPMI1640 with 10% fetal bovine serum (Atlanta Biologicals, Flowery Branch, GA), 2 mM glutamine, penicillin (100 U/ml) and streptomycin (100 μg/ml) (Thermo Fisher Scientific, Waltham, MA). NK-92 cells were cultured in ATCC formulated medium (Alpha Minimum Essential medium without ribonucleosides and deoxyribonucleosides, with 2 mM L-glutamine, 1.5 g/L sodium bicarbonate, 0.2 mM inositol, 0.1 mM 2-mercaptoethanol, 0.02 mM folic acid, 12.5% horse serum and 12.5% fetal bovine serum) and 15 ng/ml human recombinant interleukin 2 (IL-2, R &D Systems).

### RNA interference

Silencer Select siRNA negative control (4390844) and those targeting GFAT1 (human, s5708; mouse, s66610) or GFAT2 (human, s19305) were purchased from Thermo Fisher Scientific. siRNA was transfected with INTERFERin (PolyPlus-transfection, New York, NY) at 10 nM according to the instruction from the manufacturer.

### Coculture and enzyme-linked immunoabsorbing assay (ELISA) of IL-2

Cancer cells were seeded at a density of 50 000 cells/well in 24-well plate in triplicate overnight with 0.5 ml medium. Cells were then transfected with control or GFAT1 siRNA. Twenty-four hours post-transfection, cells were incubated with fresh medium with IFNγ (10 ng/ml) for about 6 h. Jurkat T cells were collected and resuspended in medium with 20 ng/ml PMA and 200 ug/ml ionomycin and added to cancer cells according to the ratios indicated in the figure legends. IL-2 concentration in culture medium released by Jurkat T cells was determined by an ELISA kit from Thermo-Fisher (Cat. 88-7025-88) according to the instruction of the manufacturer. T cells were then washed off and cancer cells were lysed to confirm the knockdown of GFAT1 and its effect on PD-L1 expression with Western blot.

### NK cell cytotoxicity assay

Cancer cells were seeded at a density of 20 000 cells/well in a 48-well plate in triplicate, transfected with control or GFAT1 siRNA for 24 h, or treated with DON (10 μM) for 30 min, and then incubated with IFNγ (10 ng/ml) overnight. The media were changed and NK-92 cells were added to the monolayer culture at the indicated ratios. After 4 h of coculture, the media were refreshed to remove NK cells. Cell viability was carried out after 24 h as described below. The results were expressed as relative cell viability with corresponding control set as 1.

### Cell viability assay

Cells were incubated with 40 μg/ml MTT for 2 h. After three washes with phosphate-buffered saline, DMSO was added to the wells. The color intensity (OD570) of the solutions was determined with a plate reader. Cell viability was expressed as percentages with the reading of the control set as 100. In some experiments, relative cell viability was further calculated as indicated in individual figure legends.

### Reverse transcription and polymerase chain reaction (PCR)

Total RNA of cell lines was extracted using RNeasy RNA extraction kit from Qiagen, (Valencia, CA). Reverse transcription was carried out with GoScript Reverse Transcription System (A5001; Promega, Madison, WI). The primers with the following sequences were synthesized by Integrated DNA Technologies (San Diego, CA): PD-L1, 5’- -3’ (forward) and 5’- -3’ (reverse); and β-actin, 5’-CCA GCC TTC CTT CCT GGG CAT-3’ (forward) and 5’-AGG AGC AAT GAT CTT GAT CTT CAT T-3’ (reverse). The amplification conditions were: 95°C, 20 sec; 58°C, 20 sec; 72 °C, 20 sec for 28 cycles for PD-L1 and 20 cycles for β-actin. The PCR products were run in 2.5% agarose with 0.5 μg/ml ethidium bromide, visualized and photographed using Bio-Rad ChemiDoc Imaging System (Hercules, CA).

### Western blot and immunoprecipitation

Cells were lysed in M2 buffer (20 mM Tris-HCl, pH7.6; 0.5% NP-40, 250 mM NaCl, 3 mM ethylene glycol-bis(aminoethyl ether)-tetraacetic acid, 3 mM ethylenediaminetetraacetic acid, 2 mM dithiothreitol, 0.5 mM phenylmethylsulfonyl fluoride, 20 mM β-glycerophosphate, 1 mM sodium vanadate and 1 μg/ml leupeptin) and subjected to a round of freeze-thaw cycle to obtained whole cell lysate. Protein concentration was determined with Bio-Rad protein quantification reagent (Cat. 500-0006). For immunoprecipitation, cell lysate containing about 500 μg protein was cleared with 30 μl Protein A (Millipore, Billerica, MA) for 4 h, then washed three times with M2 buffer prior to incubation with anti-PD-L1 antibody and 25 ul Protein A agarose overnight. The beads were collected with centrifugation and washed seven times with M2 buffer before addition of Laemmli loading buffer. Protein samples were run on 12% sodium dodecyl sulfate - polyacrylamide gel and transferred to polyvinylidene difluoride membrane, which was blocked with 5% skim milk for 2 h at room temperature, incubated with primary antibody overnight at 4 °C and then with secondary antibodies at room temperature for 1 h. Signal was developed with chemiluminescence reagent (Millipore), visualized and recorded using Bio-Rad ChemiDot Imaging system.

### Statistics

Quantitative data were expressed as mean ± SD (standard deviation). Statistics was done with GraphPad Prism 5.0 software using two-tailed unpaired *t* test to compare two means. P<0.05 was considered statistically significant.

## Results

### IFNγ induces PD-L1 expression in lung cancer cells without activating HBP

The effects of IFNγ stimulation on the expression of PD-L1 and the activation of the HBP were investigated. Consistent with previous studies(11,12), IFNγ increased PD-L1 expression in human lung adenocarcinoma cell lines regardless of common gene mutation status (wild type or mutant p53, K-ras or EGFR) and basal levels of the protein, and in murine Lewis lung cancer cells (LLC1). All human lung cancer cell lines expressed GFAT isozymes (GFAT1 and GFAT2) at various abundances. LLC1 expressed GFAT1, but GFAT2 was undetectable in the cells. No induction of the two enzymes by IFNγ was observed. OGT, the enzyme that catalyzes the attachment of O-GlcNAc, also did not change after IFNγ stimulation. Accordingly, protein O-GlcNAcylation levels indicative of HBP activity remained largely the same in treated and control cells (Figure 1A and 1B; Supplemental figure 1).

**Figure 1.**
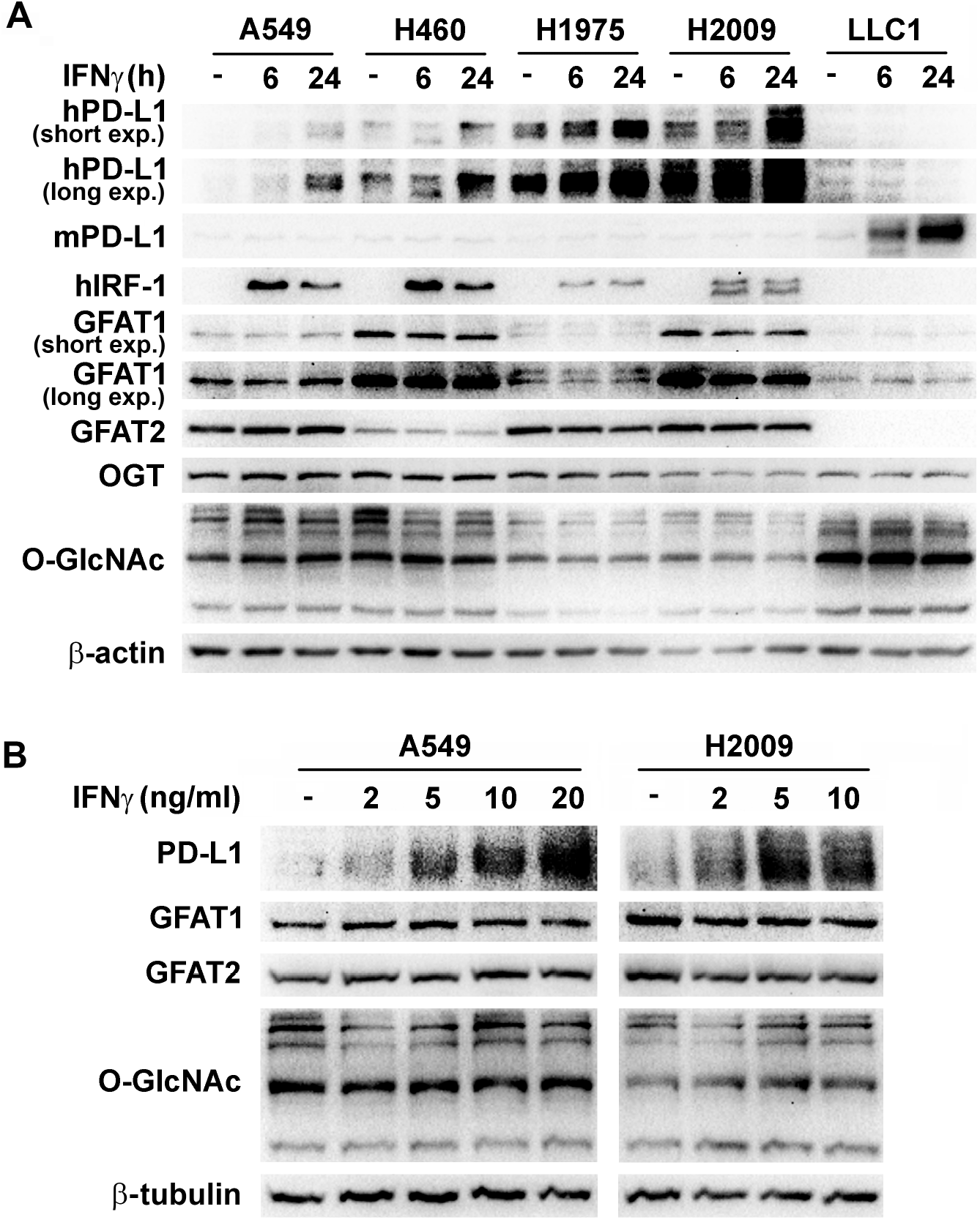
IFNγ induced PD-L1 expression in lung cancer cells without activation of HBP. (**A**) Lung cancer cell lines were treated with 10 ng/ml IFNγ for 4 h, 6 h and 24 h. (**B**) H2009 and A549 cells were treated with increasing concentrations of IFNγ for 24 h. Protein levels were determined by Western blot in total cell lysates with respective antibodies. β-Actin or β-tubulin was used as a loading control.

### Basal GFAT1 activity is indispensable for IFNγ induced PD-L1 expression

Because lung cancer cells generally have higher HBP activity in comparison with normal bronchial epithelial cells(13), how basal GFAT activity affects IFNγ-induced PD-L1 expression was examined. SiRNA knockdown of GFAT1 in all the human lung cancer cell lines tested (A549, H2009, H1975, SK-MES-1, and SW900) markedly reduced PD-L1 levels induced by IFNγ, while GFAT2 knockdown exerted minimal effects (Figure 2A, Supplemental Figure 2A, and not shown). In LLC1 cells with undetectable GFAT2, IFNγ increased the expression of PD-L1 and the induction was suppressed by knocking down GFAT1 (Figure 1A and Supplemental Figure 2B). Further, DON as well as azaserine, both glutamine analogs and inhibitors of GFAT, recapitulated the effect of GFAT1 siRNA in suppression of IFNγ-induced PD-L1 expression in cancer cells (Figure 2B and Supplemental Figure 2B). In contrast, CB-839, which inhibits glutaminase, also a target of glutamine analogs, did not reduce PD-L1 expression (not shown). These results suggest IFNγ-induced PD-L1 expression depends on GFAT1 but not GFAT2.

**Figure 2.**
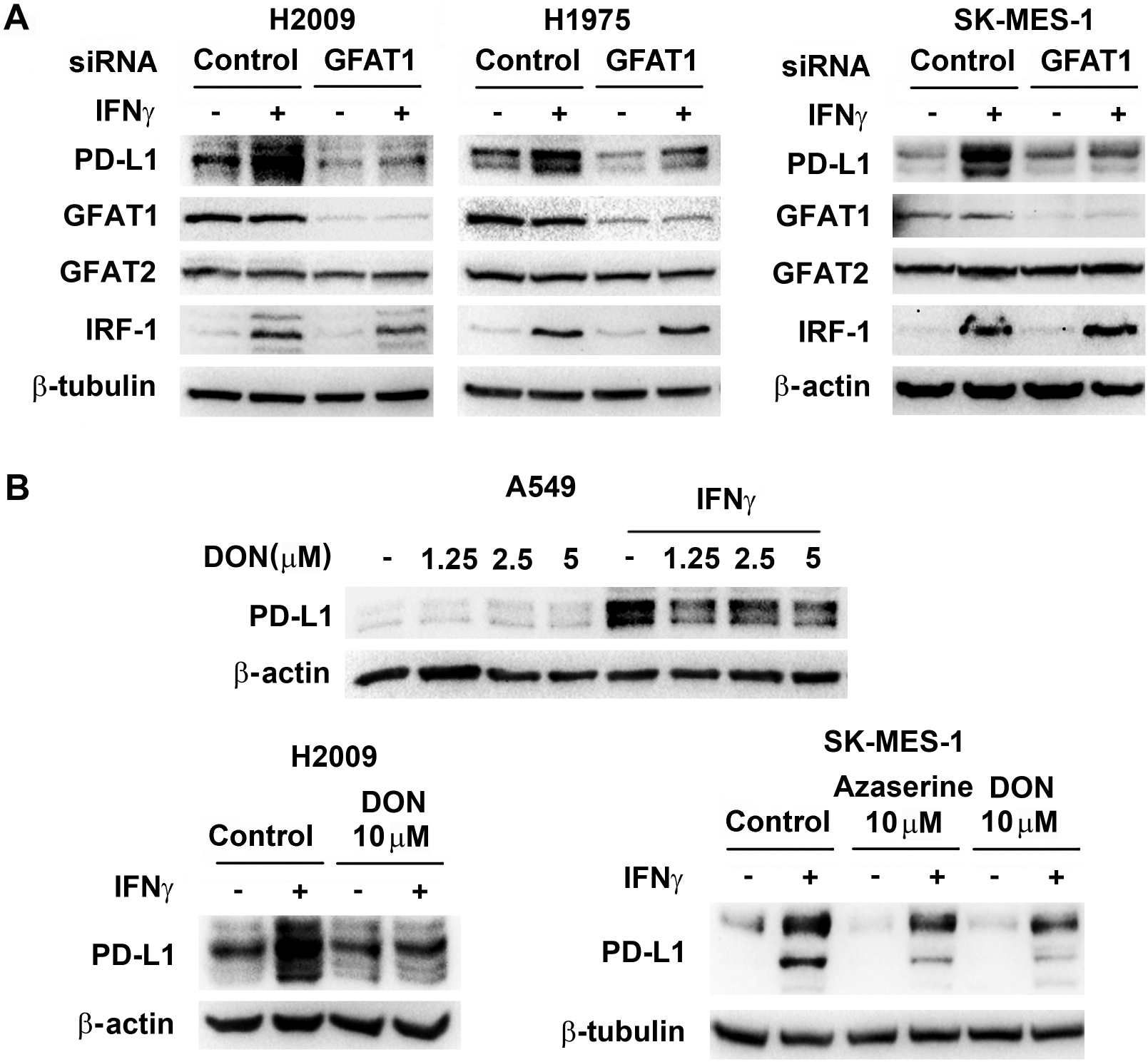
Knockdown of GFAT1 or inhibition of its activity decreases IFNγ-induced PD-L1 protein levels in cancer cells. Cancer cells were transfected with control or GFAT1 siRNA for 24 h (A), or incubated with azaserine (10 μM) or DON at the indicated concentrations for 30 min (B), prior to treatment with IFNγ (10 ng/ml) for another 24 h. Western blot was carried out to detect the expression of proteins in total cell lysates using β-actin or β-tubulin as a loading control.

### GFAT1 affects PD-L1 protein stability

The mechanism of suppression of PD-L1 expression by inhibiting GFAT1 was investigated. Phosphorylation of STAT1 and STAT3 and the induction of IRF-1 (target gene of STAT1) by IFNγ (14) were comparable in GFAT1 knockdown cells and control cells (Figure 2A and not shown). Furthermore, GFAT1 knockdown did not decrease IFNγ-induced PD-L1 mRNA expression in various cancer cells (Supplemental Figure 3). These results suggest that GFAT1 does not directly modulate IFNγ signaling. In addition, GFAT1 knockdown did not impede the cycling of PD-L1(15) as assessed by treating cells with brefeldin A, which interrupts the transport of protein from endoplasmic reticulum to Golgi apparatus. Treatment of knockdown cells with inhibitors that suppress autophagy and lysosome activity (CQ, VSP34 inhibitor, and 3-MA) were also not able to recover PD-L1 level (Figure 3A and 3B). On the other hand, proteasome inhibitor MG132 increased basal level of PD-L1 and partially reversed the effect of GFAT1 knockdown on PD-L1 expression (Figure 3B). In GFAT1 knockdown H2009 cells, the basal PD-L1 level was lower (Figure 2A, 3A and 3C), and there was a 4-fold increase in the rate of degradation of the protein (half-life 4.8 h compared to over 22 h projected in control cells; Figure 3C and 3D). Therefore, inhibition of GFAT1 accelerated degradation of PD-L1 through the proteasome pathway (Figure 3B)(12).

**Figure 3.**
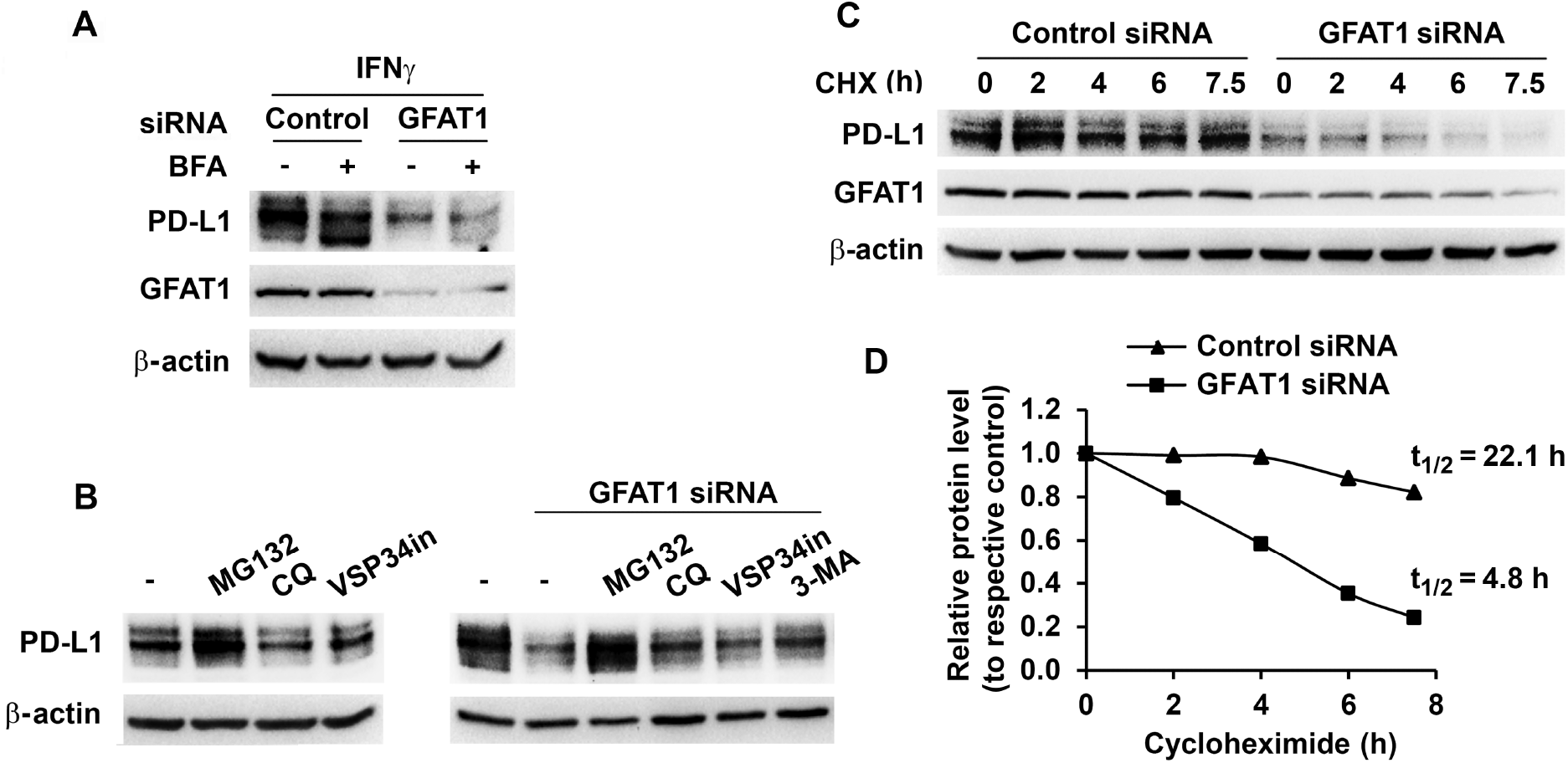
Knockdown of GFAT1 accelerates PD-L1 degradation by proteasomal pathway. (**A**) H2009 cells were transfected with control or GFAT1 siRNA for 24 h, and then incubated with IFNγ for 6 h with or without prior brefeldin A (BFA, 5 μM) treatment for 30 min. (**B**) H2009 cells were treated with inhibitors (MG132, 1 μM; CQ, 20 μM; and 3-MA, 5 mM; left panel); or transfected with control or GFAT1 siRNA. Six hours posttransfection, media were refreshed and inhibitors (MG132, 1 μM; CQ, 20 μM; VSP34 inhibitor, 10 μM; and 3-MA, 5 mM) added. Total cell lysate was prepared 24 h later for Western blot. Low concentration (1 μM) of MG132 was used to minimize cell death in this setting. **(C** and **D)** H2009 cells were transfected with control or GFAT1 siRNA for 24 h. Cycloheximide (CHX, 10 μM) was added to the cells at different time points. PD-L1 level were detected by Western blot. Band densities were quantified and normalized to loading control and the half-life of PD-L1 was calculated with corresponding untreated control set as 1.

### GFAT1 activity is important for glycosylation of PD-L1

Recent studies indicate that N-linked glycosylation is essential for PD-L1 protein stability(12). The molecular weight of native PD-L1 is about 37 kDa. Glycosylation results in species with higher molecular weights that accumulate at about 50 kDa. Treatment with tunicamycin, which inhibits N-linked glycosylation, strongly reduced the total induced PD-L1 protein with the presence of lower molecular weight bands representing unglycosylated and lower glycosylated species of PD-L1 (Figure 4A). Since HBP product GlcNAc is the major building block for protein glycosylation, we investigated if inhibition of GFAT1 could interfere the glycosylation of PD-L1. Similar to results from TM treatment, administration of high concentration of DON decreased total PD-L1 with resultant low molecular weight proteins (Figure 4B). In GFAT1 knockdown cells, there was an increase of PD-L1 species with molecular weight around 37 kDa (Figure 4C), suggesting newly synthesized protein had less glycosylation.

**Figure 4.**
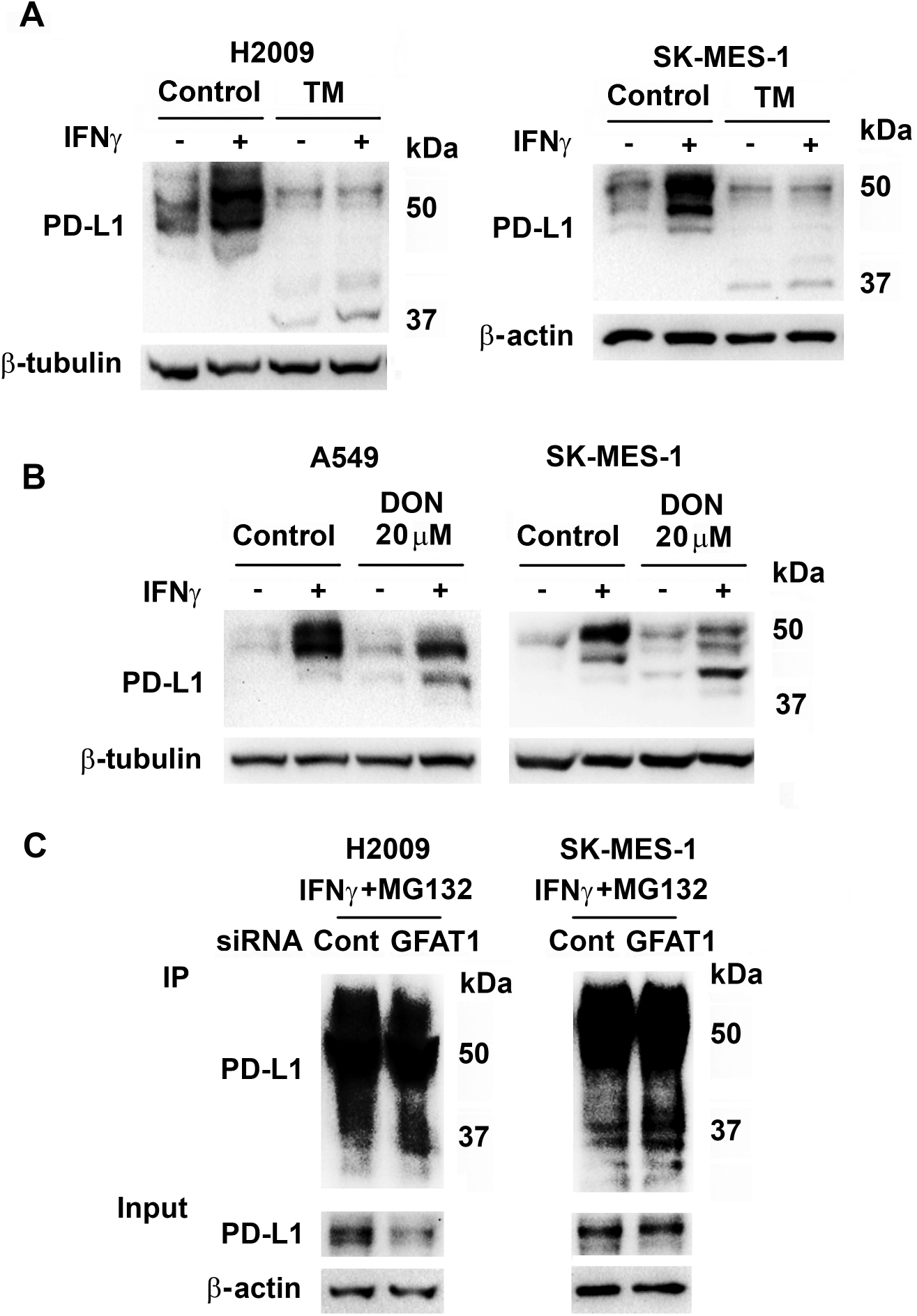
Knockdown of GFAT1 decreases glycosylation of PD-L1. (**A**) H2009 and SK-MES-1 cells were treated with tunicamycin (TM, 2 μM) 30 min before addition of IFNγ for 24 h. (**B**) A549 and SK-MES-1 cells were treated with DON (20 μM) 30 min prior to addition of IFNγ for 24 h. Protein was detected in total cell lysates with Western Blot (**A** and **B**). (**C**) H2009 and SK-MES-1 cells were transfected with control or GFAT1 siRNA for 24 h. The cells were incubated with medium with IFNγ for 6 h before the addition of MG132 (1 μM for H2009 and 2 μM for SK-MES-1). The next day, cells were lysed for immunoprecipitation with PD-L1 antibody (Cell Signaling, 7 μl used for H2009 and 5 μl used for SK-MES-1) and Protein A agarose. Western blot was carried out with 5% lysate as inputs using TrueBlot rabbit second antibody.

### Inhibition of GFAT1 in lung cancer cells diminishes the suppression of T cell activation and enhances NK cells killing of IFNγ-treated cancer cells

The functional consequence of inhibition of GFAT1 activity and PD-L1 expression was assessed by knocking down GFAT1 in cancer cells, or treating cancer cells with DON before exposure to IFNγ followed by culturing the treated cancer cells with activated T cells or NK cells. Compared to those cocultured with control cancer cells, activated T cells released about 50% more IL-2 in the presence of cancer cells transfected with GFAT1 siRNA or treated with DON (Figure 5A and Supplemental Figure 4). Similarly, NK cells were able to kill more cancer cells with lower GFAT activity to various extents across cell lines (Figure 5B).

**Figure 5.**
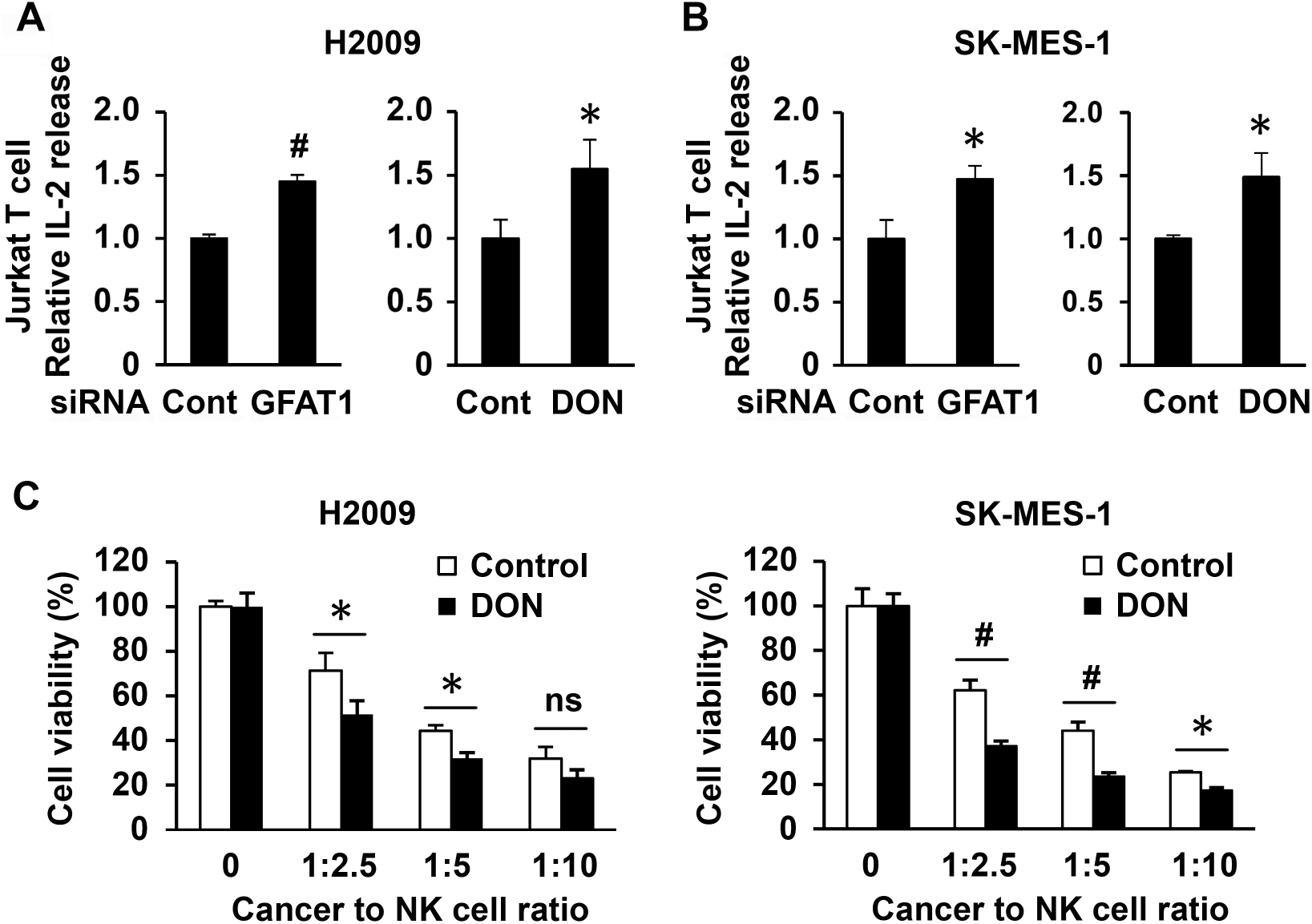
Knockdown of GFAT1 or DON treatment in cancer cells increases T cell activation and NK cell killing. (**A**) Cancer cells were seeded in triplicate in 24-well plates at a density of 5 × 10^4^ cells/well. The next day, cells were transfected with control or GFAT1 siRNA for another 24 h. The cells were then incubated with media containing IFNγ (10 ng/ml) for 6 h. The medium was then removed and Jurkat T cells in medium with 25 ng/ml PMA and 200 ug/ml ionomycin were added to cancer cells. The ratio of cancer cell to T cell was 1:2 for H2009 and 1:4 for SK-MES-1. The next day (18 h for H2009 and 24 h for SK-MES-1), the media were collected for ELISA for IL-2 concentration. For DON inhibition, 10 μM DON was added overnight before the treatment of IFNγ, the ratio of cancer cell to T cell was 1:3 and incubation time was 24 h for both cell lines (**B**). (**C**) Cancer cells were seeded and transfected with siRNAs and treated with IFNγ as described in (A). NK cells were added to cancer cells in the indicated ratio for 4 h. The media were refreshed and NK cells removed. MTT assay was carried out 24 h later. *, P<0.05; #, P<0.01; ns, not significant.

## Discussion

Currently PD-L1 is targeted by monoclonal antibodies in clinical practice. Other compounds that can modulate PD-L1 expression could provide alternatives to antibodies with different pharmacological properties and toxicity profiles, which may be used in combinational therapy. For example, metformin induces degradation of PD-L1 through endoplasmic reticulum-associated (16) or other signaling pathways(17), and was shown to improve overall response and disease control rate in combination with checkpoint inhibitors in small clinical trials in NSCLC(18) and melanoma(19). Many natural compounds (e.g. Triptolide(20), Galic acid(21) and Resveratrol(22)) were also able to inhibit the expression of PD-L1 in cancer cells through various mechanisms. Like metformin, these compounds are multifunctional rather than just affecting the expression of PD-L1. There are also efforts to screen small molecules(23), peptides(24) and kinase inhibitors(25) for PD-L1 inhibitors.

Modulating turnover is a novel approach to control PD-L1 protein level. PD-L1 is heavily N-glycosylated which increases its stabilization. Several proteins are involved in posttranslational modification, trafficking and degradation of PD-L1. Protease COP9 signalosome 5 (CSN5) was shown to inhibit the ubiquitination and proteasomal degradation of PD-L1(26). Further, CKLF-like MARVEL transmembrane domain-containing protein (CMTM) 4 and CMTM 6 are proteins maintaining the cell surface expression of PD-L1 and reducing its endocytosis and degradation in lysosome(15,27). Targeting these proteins as well as glycosylation to increase the degradation of PD-L1 is in development.

This work provides compelling evidence for the regulation of PD-L1 by GFAT1, a rate-limiting enzyme of the hexosamine biosynthesis pathway (HBP). We show that inhibition of GFAT1 activity, and probably other enzymes of HBP, can increase the degradation of PD-L1 and thus alleviate the inhibitory effects of cancer cells on the function of T cells and NK cells. The increasing degradation of the protein is attributed to decreased glycosylation when GFAT1 activity is impaired, although other mechanisms cannot be ruled out. The advantage of inhibiting GFAT1 is that DON is available as a prototypic GFAT inhibitor and a research tool. As a glutamine analog, DON is not specific to GFAT, but it has gone through numerous clinical trials in the past, therefore its toxicity is known and efforts to minimize the side effects are ongoing(28,29). New inhibitors for GFAT are also in development(30–32). Another advantage of targeting GFAT/HBP is that inhibition of the pathway itself has anti-cancer properties, as we (13)and others have demonstrated(33–36). HBP is frequently hyperactive in cancer cells. Inhibition of HBP can thus have much broader influences besides PD-L1 degradation in cancer cells in systemic therapy, which may cause side effects but increase efficacy against cancer. Therefore, our study supports developing therapeutics to modulate HBP/GFAT1 activity to augment the effectiveness of immunotherapy using checkpoint blockers.

## Abbreviations

BFA: Brefeldin A
CMTM: CKLF-like MARVEL transmembrane domain-containing protein
CQ: chloroquine
CSN5: protease COP9 signalosome 5
CTLA-4: cytotoxic T lymphocyte antigen 4
DMSO: dimethyl sulfoxide
DON: 6-diazo-5-oxo-L-norleucine
EGFR: epidermal growth factor receptor
GFAT: L-glutamine:D-fructose-6-phosphate amidotransferase
GlcNAc: N-acetyl-glucosamine
GlcN: glucosamine
GLS: glutamine synthase
HBP: hexosamine biosynthesis pathway
IFNγ: interferon gamma
IRF-1: interferon gamma response factor 1
3-MA: 3-methyladenine
NSCLC: non-small cell lung cancer
OGT: O-GlcNAc transferase
PD-1: programmed death 1
PD-L1: programmed death ligand 1
TM: tunicamycin
UDP: uridine diphosphate

## Acknowledgements

This paper is dedicated to Dr. Yong Lin, a highly valued member of the Lovelace Lung Cancer Program who lost his battle to cancer in April 2020. Yong was a productive molecular biologist who made many important contributions toward understanding the mechanisms of programmed cell death and deregulation of signal transduction pathways in cancer.

## Data availability statement

The data that support the findings of this study are available from the corresponding author upon reasonable request.

## Supplemental figures

**Figure S1.**
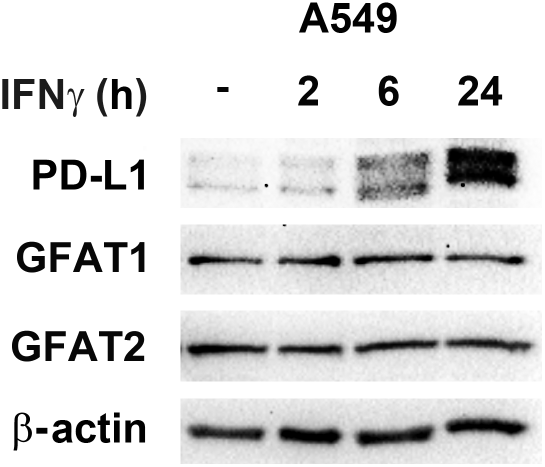
Time-dependent induction of PD-L1 but not GFAT by IFNγ. A549 cells were treated with 10 ng/ml IFNγ for various times. Protein levels were detected with Western Blot using respective antibodies with β-actin as a loading control.

**Figure S2.**
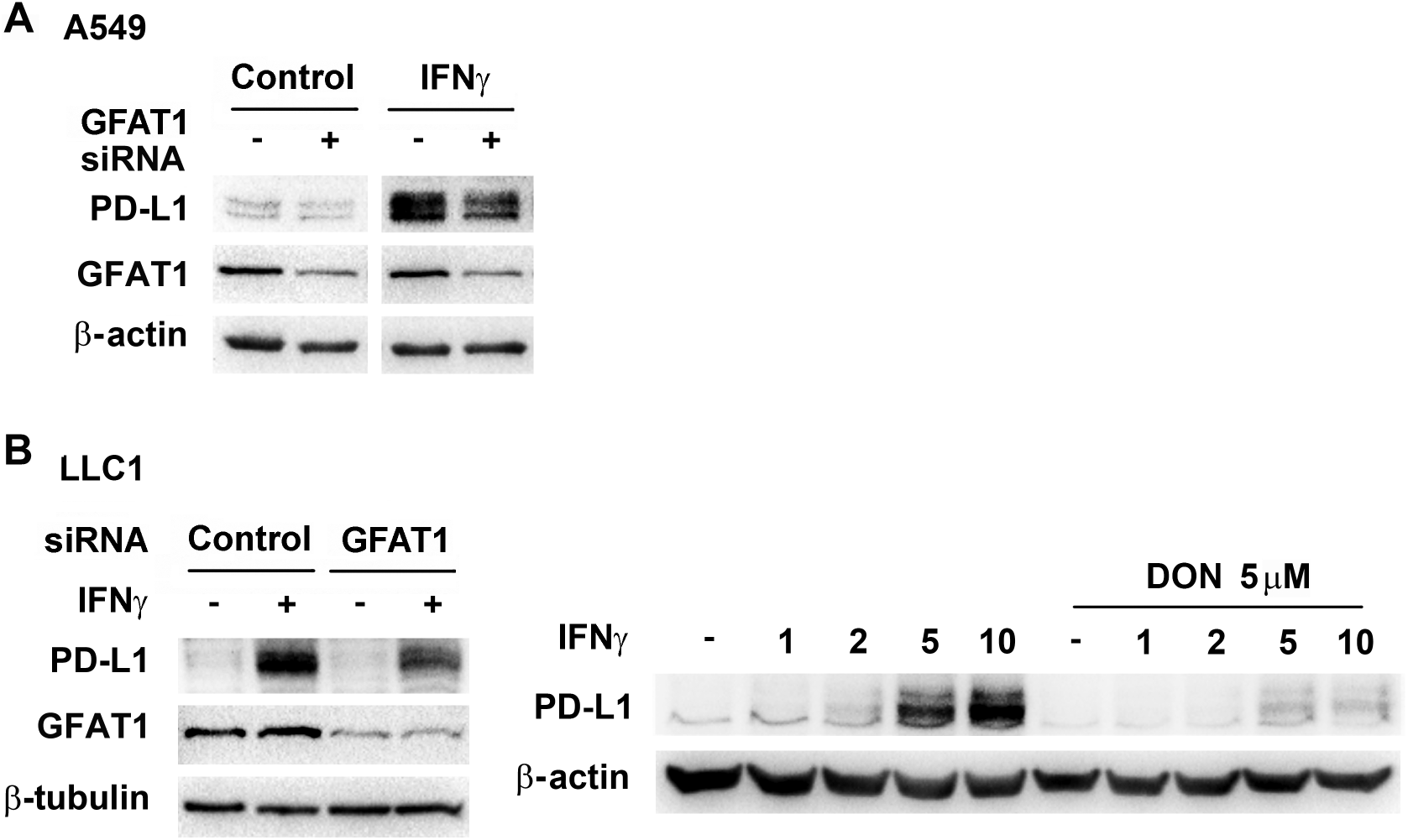
Inhibition of GFAT1 activity suppresses IFNγ-induced PD-L1 expression. (**A**) A549 cells transfected with control or GFAT1 siRNA for 24 h, then treated with 10 ng/ml IFNγ for 24 h. (**B**) LLC1 cells were transfected with siRNAs for 24 h, or treated with DON (5 μM) for 30 min, prior to the incubation of 10 ng/ml IFNγ or various concentrations of the cytokines, for 24 h. Protein levels were detected with Western Blot using respective antibodies with β-actin as a loading control. Western Blot was used to detect the proteins with loading controls.

**Figure S3.**
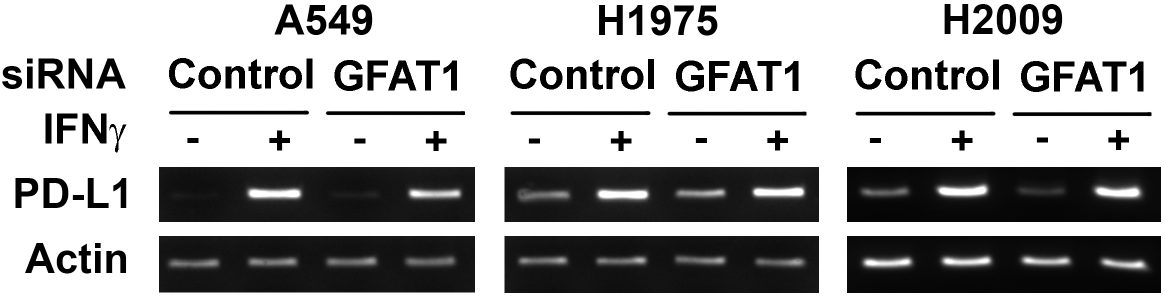
Knockdown of GFAT1 does not affect IFNγ-induced transcription of PD-L1. Lung cancer cell lines were transfected with control or GFAT1 siRNA for 24 h and then left untreated or treated with IFNγ for 24 h. Cells were harvested for mRNA extraction and RT-PCR for PD-L1 and β-actin as a loading control.

**Figure S4.**
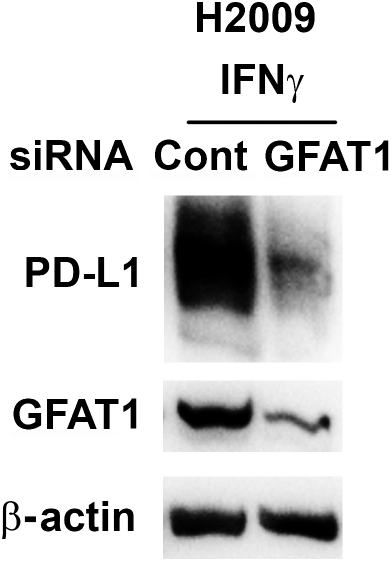
GFAT1 knockdown cells maintains lower levels of PD-L1 in the presence of Jurkat T cells in medium containing PMA and ionomycin. H2009 cells were transfected with siRNAs and treated with IFNγ as previously described. Cells were then incubated with Jurkat T cells in activation medium for 24 h. After cultured media were removed for IL-2 detection, cells in wells were washed, lysed and combined for Western Blot. β-Actin served as a loading control. Cont, control.

